# The effect of repeated periods of drought and aestivation on *Allolobophora chlorotica* reproductive output

**DOI:** 10.1101/2025.11.04.686498

**Authors:** R. V. A. Bray, M. E. Hodson, P. J. Watt

## Abstract

Increasing drought frequency under climate change is expected to intensify periods of suboptimal soil moisture, adversely affecting earthworms and other soil biota. To survive desiccation, some earthworms enter aestivation, a state of reduced metabolic activity during which reproduction and ecosystem service provision are suspended. However, the capacity of earthworms to recover from repeated drying events remains poorly understood. This study examined how multiple drought-aestivation cycles influence the reproductive performance of *Allolobophora chlorotica*, one of the most common UK earthworm species. Adults were exposed to one, two or three 14-day drying periods to gravimetric moisture contents of ∼11-13 wt% in loamy soil, each followed by three days under favourable moisture conditions. After the final exposure, earthworms were placed in groups of four into optimally moist soil for 50 days to reproduce. Cocoon number, mass, viability and incubation time were measured as indicators of reproductive success. Cocoon production declined significantly with increasing aestivation frequency (p < 0.001), being lowest during Days 1-10 of recovery (∼0-0.02 cocoons earthworm^-1^ day^-1^), peaking between Days 20-30 (∼0.08-0.16 cocoons earthworm^-1^ day^-1^) and declining over the final 20 days (∼0.08-0.12 cocoons earthworm^-1^ day^-1^). Unexpectedly, earthworms previously exposed to drying and aestivation produced more (226 vs 171 cocoons, p < 0.05) and heavier (6.701 ± 1.205 vs 6.036 ± 1.256 mg, p < 0.05) cocoons than those kept under constant high moisture, suggesting compensatory growth upon rehydration and possibly reflecting food limitation or soil compaction in controls. Cocoon viability and incubation time did not differ significantly between treatments. Across treatments, earthworm mass strongly predicted fecundity as heavier individuals produced more (p < 0.001) and heavier (p <0.001) cocoons, and cocoon mass was positively correlated with hatching success (p < 0.001). Overall, *Al. chlorotica* displayed resilience to short-term, intermittent drought through aestivation, but reproductive success remained sensitive to the combined effects of soil moisture, food availability, and soil structure. These findings highlight the importance of considering multiple environmental constraints when predicting soil fauna responses to increasing drought frequency and duration.

## Introduction

The frequency and severity of climatic extremes are projected to increase under climate change (Seneviratne *et al*., 2021). Models based on intermediate greenhouse gas emission scenarios predict widespread drought and reduced soil moisture in the upper 10 cm of soil across many land areas during the 21^st^ century (Dai, 2013). Such declines in soil water availability threaten important soil organisms, including earthworms, which depend on water to facilitate oxygen uptake for respiration (Holmstrup, 2001), and to form coelomic fluid which acts as a hydrostatic skeleton, supporting body turgor and locomotion (Kretzschmar and Bruchou, 1991; Ramsay, 1949; Whalen *et al*., 2000). To survive desiccation, some earthworm species enter aestivation, coiling into a knot and entering a state of reduced metabolic activity until soil moisture improves (Gerard, 1967; Jiménez *et al*., 2000; McDaniel *et al*., 2013a; McDaniel *et al*., 2013b). Prolonged or more frequent earthworm aestivation and thus inactivity will constrain their ability to deliver key ecosystem services such as bioturbation, organic matter decomposition and nutrient cycling (Blouin *et al*., 2013). Moreover, the persistence of earthworm populations depends on their capacity to reproduce and expand spatially, both of which may be compromised by the energetic demands of repeated aestivation and the reduced opportunities for mating associated with longer and more frequent drought (McDaniel *et al*., 2013a).

Earthworms are simultaneous hermaphrodites possessing both male and female reproductive organs. During mating, individuals mutually exchange sperm and seminal fluids from the male pores to the spermathecae of their partner (Porto *et al*., 2012; Christyraj *et al*., 2025). Sperm and an egg are released into a mucus band secreted by the clitellum (Christyraj *et al*., 2025) which hardens to form an ellipsoid cocoon (Sherlock, 2018). Reproductive rate and cocoon mass vary both among earthworm species and with environmental conditions. Reduced reproductive investment can occur due to poor body condition resulting from indirect or direct effects of suboptimal conditions on growth and behaviour (Reinecke and Reinecke, 2007). Several studies demonstrate trade-offs between growth and reproduction under environmental stress. For example, *Lumbricus rubellus* reduced cocoon production in calcium-poor soil (West *et al*., 2003), and *Esenia andrei* maintained growth at the expense of reproduction under unfavourable pH and temperature conditions (soil pH > 7, at and below 15 °C, and at 30 °C) (van Gestel *et al*., 1992). Similarly, *Esenia fetida* subjected to mechanical stress produced fewer and lighter cocoons while maintaining somatic growth (Aira *et al*., 2007).

Soil water strongly influences earthworm reproduction. It can be expressed gravimetrically relative to the dry mass of soil (Weil and Brady, 2017), or in terms of water potential, which indicates the force at which water is held in the soil and thus how available it is for uptake (Kretzschmar and Bruchou, 1991). Holmstrup (2001) found *Aporrectodea caliginosa* exhibited optimal cocoon production at water potentials below ∼pF 2 (10 kPa, above ∼15-17 wt%), reduced output above ∼pF 2.08 (12 kPa, below ∼13 wt%), and complete inhibition above ∼pF 2.6 (40 kPa, below ∼9-11 wt%). Likewise, *Allolobophora chlorotica* produced the most cocoons around pF 2 (10 kPa), with production inhibited at pF 3 (100 kPa) (Evans and Guild, 1948).

Cocoon development and hatching success also depend on environmental conditions, particularly the soil moisture content (Jensen and Holmstrup, 1997). This is notable because cocoons are typically deposited in upper soil layers which are more exposed to fluctuating conditions and lose water at a faster rate than deeper layers (Nepstad *et al*., 2002). For instance, Butt (1997) found that all *Al. chlorotica* cocoons occurred within the top 10 cm of soil, with over half within the top 5 cm. Evidence suggests that once formed, earthworm cocoons are resistant to harsh conditions including desiccation, although this is dependent on the rate and extent of dehydration. For instance, *Dendrobaena octaedra* cocoons tolerated gradual losses of ∼95 % of their water content, yet rapid desiccation exposure below 89 % RH (relative humidity) caused 100 % cocoon mortality (Petersen *et al*., 2008). Such tolerance allows earthworm populations to recover following climatic perturbations once conditions improve, even if adults and juveniles perish.

Despite increasing knowledge of drought effects on cocoon production and survival, the long-term reproductive consequences of repeated drought and aestivation remain largely unexplored. Understanding these carry-over effects if critical for predicting the recovery time required for earthworm populations under future drought regimes. Estimates suggest that approximately 4-6 weeks of favourable moisture are necessary for completion of a full earthworm reproductive cycle and for hatchlings to accumulate biomass (Edwards and Bohlen, 1996; Ruiz *et al*., 2021). Consequently, droughts recurring more frequently than every 4-6 weeks may prevent full population recovery. Moreover, Holmstrup (2001) demonstrated that the negative effects of drought exposure can persist long after rewetting, with *Aporrectodea caliginosa* requiring between two weeks to two months to resume normal cocoon production following 14-day drying bouts, depending on the degree of drying experienced.

The present study examines how repeated bouts of drying and aestivation, followed by recovery under optimal moisture, affect the mass and reproductive output of *Al. chlorotica*. Specifically, we assessed whether increasing numbers of drought-aestivation cycles have lasting negative effects on cocoon production, cocoon mass, hatching success (viability) and development time. We predicted that (i) both cocoon number and mass would decline with increasing aestivation frequency, and (ii) when transferred to more favourable soil moisture conditions, reproductive output would initially be low but increase with time spent in these conditions as earthworms recovered from aestivation.

## Methods

### Earthworm collection and culturing

Adult *Al. chlorotica* (green morph) possessing a developed clitellum and tubercula pubertatis were collected in October 2024 by hand-sorting soil from pits (18 cm x 18 cm at the surface to 20 cm depth) dug in the margins of Warren Paddock, Spen farm, Tadcaster (53°52’25.9”N, 1°19’33.4”W). The site has a well-drained, loamy and calcareous soil (Holden *et al*., 2019). Earthworms were maintained at 15 ± 1 °C in complete darkness in a controlled temperature room at the University of Sheffield. Individuals were kept in field soil maintained at 25-30 wt% moisture by addition of distilled deionised water (DIW). Temperature, light and moisture conditions followed Lowe and Butt (2005). Earthworms were fed *ad libitum* with oven-dried (105 °C) sheep manure, milled to <2 mm, sourced from sheep not treated with anti-helminthic drugs.

### Initial acclimation

Eleven groups of 15 earthworms (n = 165) were randomly selected and placed in containers with ∼4 kg of Kettering loam (Boughton Loam, Kettering, UK purchased online from Agrigem). Sheep manure (0.5 g per earthworm) was mixed into the soil as a food source, and DIW was added to reach 25 wt% moisture. *Al. chlorotica* were kept in these conditions for one week based on Fründ *et al*. (2010) recommendations for acclimation. Afterwards, individual *Al. chlorotica* were removed from the soil and placed onto filter paper moistened with DIW for 24 hours to depurate, before being gently blotted dry with tissue and weighed on a balance.

### Drought exposures to induce aestivation

After fresh mass determination, individual *Al. chlorotica* were randomly assigned to one of four treatments: a control group (n = 60) maintained at constant optimal moisture, or a drought-exposed group (n = 105) subjected to either one, two or three 14-day periods of gradual substrate drying to induce aestivation (**Fig. 1**). Mean earthworm starting mass (0.287 ± 0.039 g) did not differ significantly between the drought or control treatments (ANOVA: F = 0.009, df = 1, 114, p = 0.925), or among those assigned to one, two, and three exposures (ANOVA: F = 1.118, df =1, 114, p = 0.33) (**Fig. S1**). Earthworms were housed individually in 300 ml plastic cups containing soil packed to an air-dried density of ∼1.13 g/cm^3^ (170 g of Kettering loam and 0.5 g of sheep manure milled to <2 mm). Control soils were kept at 25 wt% moisture, considered within the optimal range for *Al. chlorotica* (Lowe and Butt, 2005) and sufficient to prevent aestivation in Tilikj and Novo (2022) experiments with *Carpetania matritensis*. Drought treatments began at 20 wt% moisture to reduce the stress of transitioning from optimal conditions. Plastic cups were covered with nylon mesh and secured with an elastic band to prevent earthworms from escaping but allow evaporation. Soils in the drought-exposed treatment were left to air-dry gradually, with no addition of water to simulate natural conditions whereby desiccation stress would increase over a period of days or weeks (Petersen *et al*., 2008).

**Figure 1.**
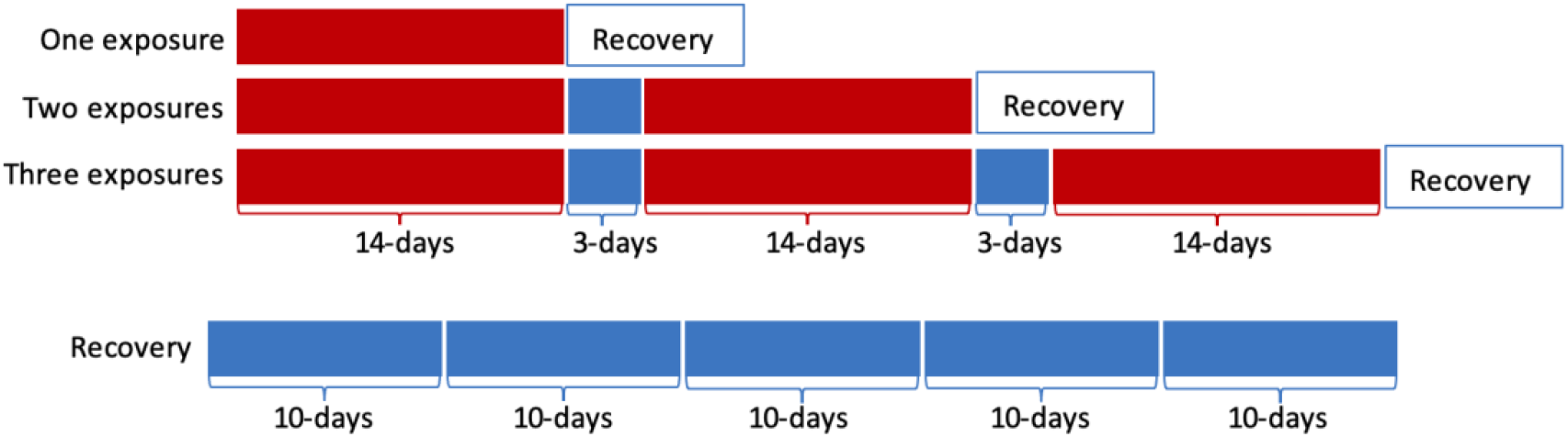
Summary of experimental design. *Al. chlorotica* were subject to 1, 2 or 3 bouts of drying (red), broken up by 3-day periods of optimal moisture conditions (blue) and a 50-day recovery period in which reproductive output was assessed every 10 days.

After 14 days, soils were destructively sampled. Individuals were classified as active (extended, mobile) or aestivating (coiled, usually within a soil chamber), then weighed on a balance. Non-aestivating individuals in the drying treatment were removed from the experiment. Of the earthworms found aestivating, 20 were randomly selected to undergo a recovery period in soil under optimal moisture conditions, while the remainder underwent a 3-day rehydration (25 wt% moisture) before transfer to fresh soils at 20 wt% moisture for a second 14-day drying cycle (**Fig. 1**). The same procedure was repeated for a third drying period. This was done to replicate conditions expected to occur in natural context whereby earthworms are subject to intermittent periods of drought and more favourable moisture conditions resulting in transitions between active and aestivating states. Control soils were maintained at ∼25 wt% through daily weighing and addition of DIW to the soil surface using a spray bottle when necessary. Control earthworms were transferred to fresh soils at the same intervals as drought-exposed individuals to ensure equal handling.

The rate of soil water loss increased slightly with each bout of air drying, resulting in a progressively lower minimum moisture content after each 14-day period, reaching 12.85 %, 11.39 % and 10.58 % after one, two and three cycles, respectively (**Fig. 2**).

**Figure 2.**
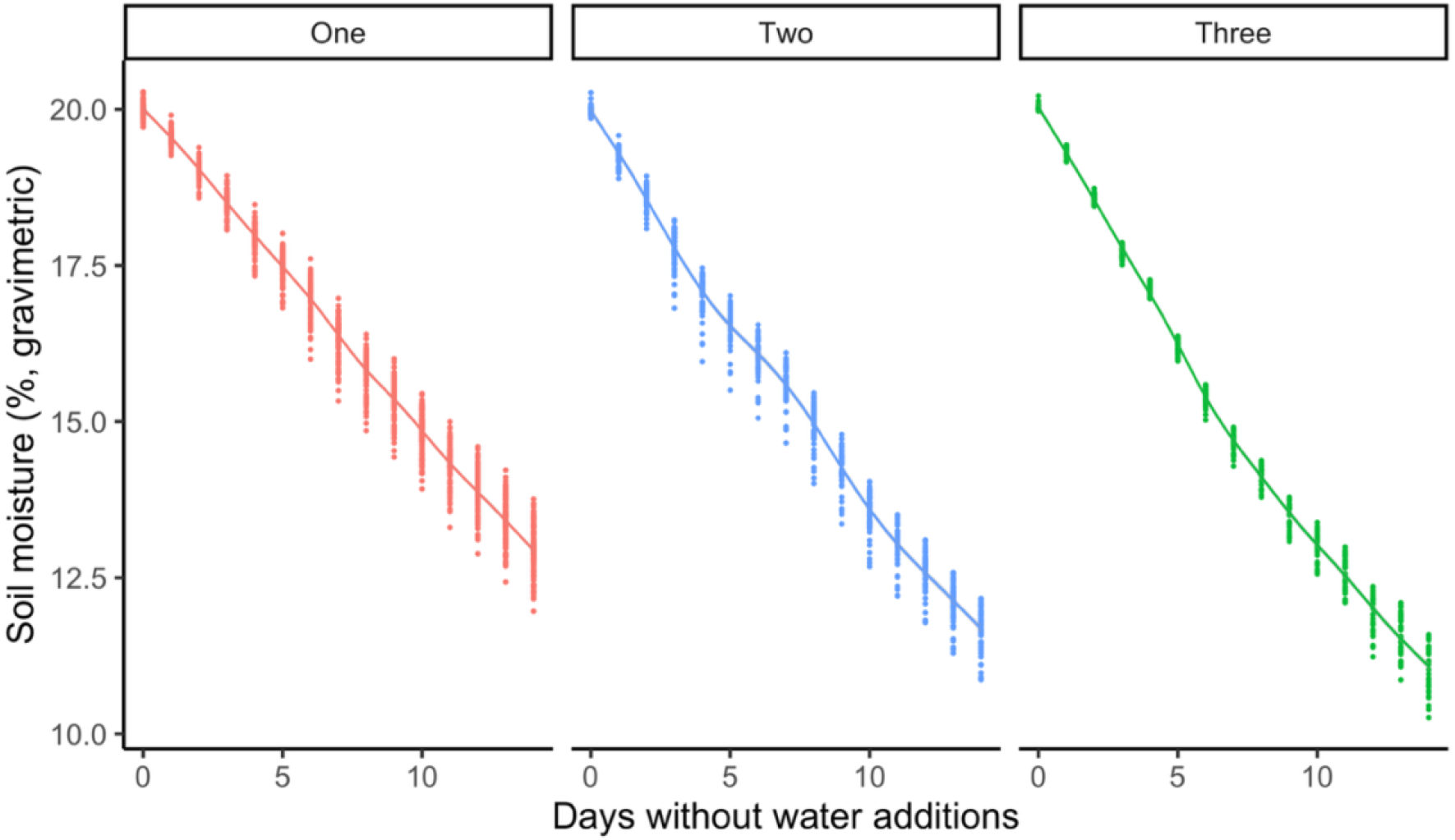
The rate of soil water loss for each of the three drying exposures. Individual data points are plotted. Red = one exposure (y = -0.51x + 20.02), blue = two exposures (y = -0.59x + 19.7) and green = three exposures (y = -0.65x + 19.67).

### Recovery period and reproductive output

Following their assigned number of drying cycles, earthworms were extracted from the soil, cleaned with a dry paper towel and weighed. Individuals were then placed onto moist filter paper for 24 hours to depurate, blotted dry and reweighed to get their fresh mass. *Al. chlorotica* were grouped by exposure history and randomly assigned in sets of four to 750 ml containers (with perforated lids) containing 400 g dry soil mixed with 2.8 g dried sheep manure (<2 mm), moistened to 25 wt% moisture with DIW. The recovery period lasted 50 days. Every 10 days, each group was extracted from the soil, cleaned with a dry paper towel, and weighed individually before being transferred together into fresh soil (**Fig. 1**). At the end of the recovery period, earthworms were again weighed pre- and post-hydration (24 hours on moist filter paper).

Soils were wet sieved through a 1 mm metal mesh to collect cocoons (Bart *et al*., 2018). Cocoons from the two- and three-exposure groups were blotted dry, weighed on a balance (to five decimal places), and then transferred to moistened filter paper in Petri dishes (Butt, 1997; Petersen *et al*., 2008). Petri dishes were incubated in a controlled temperature room under the same conditions they had been produced in (15 °C ± 1 °C, complete darkness) (**Fig. 3**). Moisture was maintained with DIW and cocoons were inspected daily for hatchlings, recording the date of hatching to determine incubation time. The experiment concluded when no new hatchlings appeared for two consecutive months.

**Figure 3.**
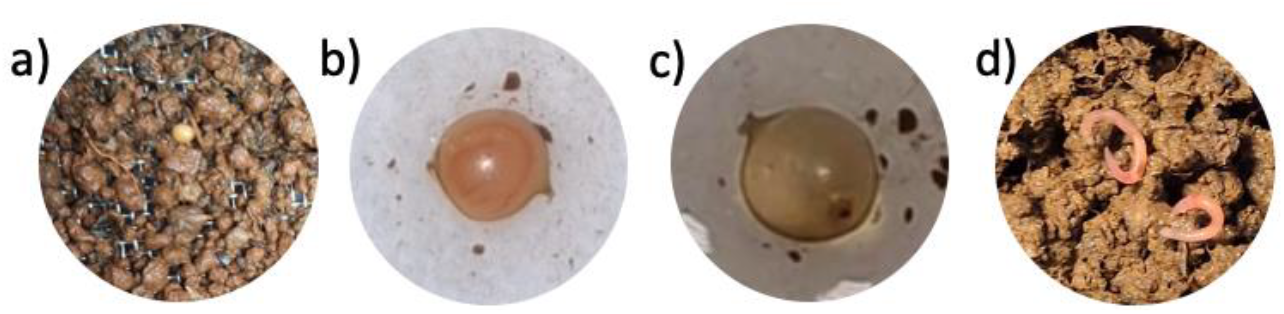
(a) A yellow *Al. chlorotica* cocoon (∼ 4 mm at longest point) seen on a 1 mm sieve after wet sieving, (b) Cocoon on moist filter paper with visible earthworm embryo developing, (c) Empty cocoon, translucent after hatching, (d) Two earthworm hatchlings (∼ 15 mm in length).

### Statistical analyses

All analyses were performed in the programming language R v4.4.1 (R Core Team, 2024). Four control individuals entered aestivation after the third 14-day period and were excluded from the analyses. No mortality occurred during the initial exposures to drying or control conditions, but three individuals died during the 50-day recovery period of optimal conditions (one from the group exposed to two bouts of drying on Day 50, and two from the group exposed to three bouts of control conditions on Day 40 and Day 50) and were excluded from cocoon production analyses. Regression of the clitellum was observed among some individuals during the experiment and so cocoon production was expressed as cocoons worm^-1^ day^-1^ per clitellate individual for each 10-day interval. Cocoon mass and hatching success were averaged per replicate group. After confirming that the assumptions of linearity, normality and equal variance were met, ANOVAs were conducted to compare the changes in mass and mean rate of cocoon production for earthworms exposed to one, two and three drying periods. Post-hoc treatment contrasts were carried out using Tukey HSD tests to identify where statistical differences occurred. Non-parametric Spearman’s rank tests were used to assess linear relationships between earthworm mass, cocoon production, mass and hatching success. Hatching success data did not meet the assumption of equal variance, so an arcsine square root transformation was first applied before a linear regression and ANOVA were carried out.

Differences in incubation time among cocoons produced during successive recovery intervals were also analysed by ANOVA.

## Results

### Changes in mass prior to reproduction

After 14 days, *Al. chlorotica* exposed to drying had lost in mass relative to their baseline initial mass, whereas individuals kept under constant optimal moisture had gained mass (**Fig. 4**). This difference was significant for earthworms exposed to a single bout of drying versus those in the constant control conditions (ANOVA: F = 35.594, df = 1, 16, p < 0.01). Time in the experiment significantly influenced mass for two- (ANOVA: F = 6.187, df = 1, 36, p < 0.05) and three-bout treatments (ANOVA: F = 19.271, df = 1, 50, p < 0.01). During 3-day recovery phases, the mass of those in the drying groups increased by ∼22-24 %, but subsequent drying losses exceeded gains, producing cumulative mass decline with each bout. In contrast, controls initially gained mass during the first 14 days, but subsequently lost mass.

**Figure 4.**
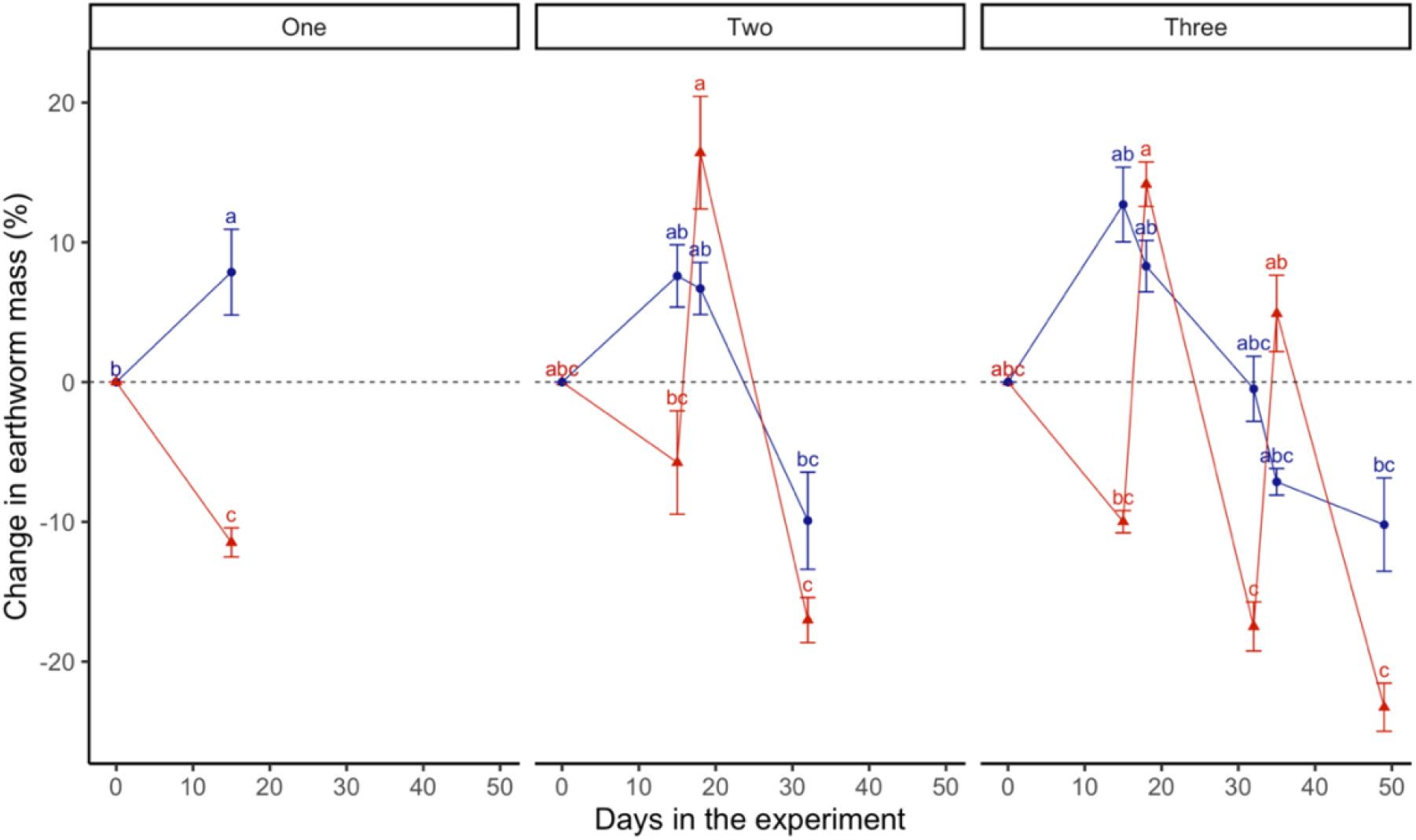
The change in mean (n = 20) earthworm mass relative to their starting mass, measured after each 14-day period of drying (red triangles) or constant moisture (blue circles) conditions and subsequent 3-day period of optimal moisture conditions. Treatments with the same letter do not differ to a statistically significant degree (p > 0.05).

After the second 14-day period, their masses either returned to their baseline starting mass or dropped below it.

### Cocoon production

Overall, total cocoon production was significantly influenced by the number of drying (or constant control) bouts experienced by *Al. chlorotica* (ANOVA: F = 12.874, df = 2, 139, p < 0.001, **Fig. S2**). Earthworms exposed to a single bout produced significantly more cocoons than those exposed to three (Tukey, p < 0.5). Earthworms subjected to two bouts also produced fewer cocoons than those with only a single exposure, but this difference was significant only under constant control conditions (Tukey, p < 0.5, **Fig. S2**).

Temporal patterns of cocoon production also differed among treatments. Between Days 10-20, significantly more cocoons were produced by earthworms that had been exposed to one drying bout compared to three (Tukey, p < 0.05, **Fig. 5**). Moreover, the duration *Al. chlorotica* spent in the recovery period also had a significant effect on cocoon production (ANOVA: F = 23.945, df = 1, 133, p < 0.001). As expected, cocoon production was initially low (Days 1-10) following aestivation but increased thereafter, peaking at Days 20-30 (**Fig. 5**). Relative to Days 1-10, cocoon production was significantly higher in Days 10-20 for earthworms exposed to a single bout of drying and in Days 20-30 for those exposed to two bouts. In contrast, earthworms exposed to three drying periods consistently produced fewer cocoons, with no significant differences across sampling intervals. After ∼30 days in the recovery period, cocoon production generally declined, although rates remained higher than during the initial 10 days for most treatment groups. Across all treatments, earthworms subjected to drying produced significantly more cocoons than those from the constant control conditions (ANOVA: F = 4.484, df = 1, 133, p < 0.05). These treatment differences were most evident in earthworms that had been subjected to multiple drying or control periods, though not statistically significant within each sampling point (Tukey, p > 0.05).

**Figure 5.**
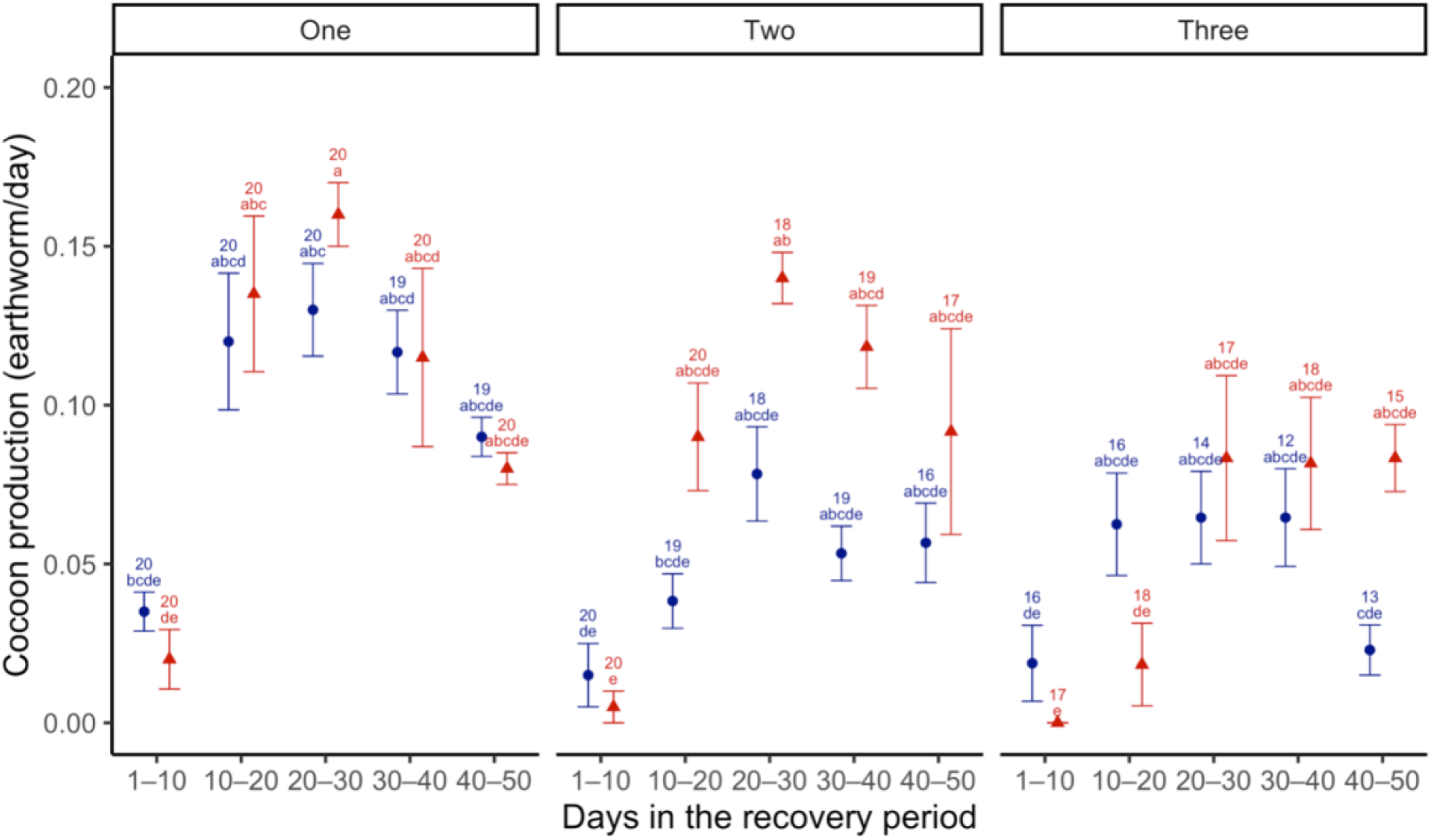
The mean cocoon production of clitellate earthworms recorded every 10 days in the recovery period from those that had been exposed to either one, two or three bouts of drying (red triangles, n = 5) or constant control conditions (blue circles, n = 4). Numbers represent live clitellate earthworms. Treatments with the same letter do not differ to a statistically significant degree (p > 0.05).

### Cocoon mass

Overall, cocoon mass was significantly greater in drought-exposed than control earthworms (ANOVA: F = 8.54, df = 1, 68, p < 0.05). Like rates of cocoon production, cocoon mass was lowest during the first 10 days of the reproductive period and increased thereafter, reaching maximum size between Days 20-40. However, these temporal differences were not statistically significant (ANOVA: F = 0.27, df = 1, 68, p = 0.605) (**Fig. 6**). Cocoon mass did not differ significantly between earthworms that had previously experienced two or three periods of drying (or the equivalent duration of constant control conditions) (ANOVA: F = 0.0004, df = 1, 68, p = 0.984).

**Figure 6.**
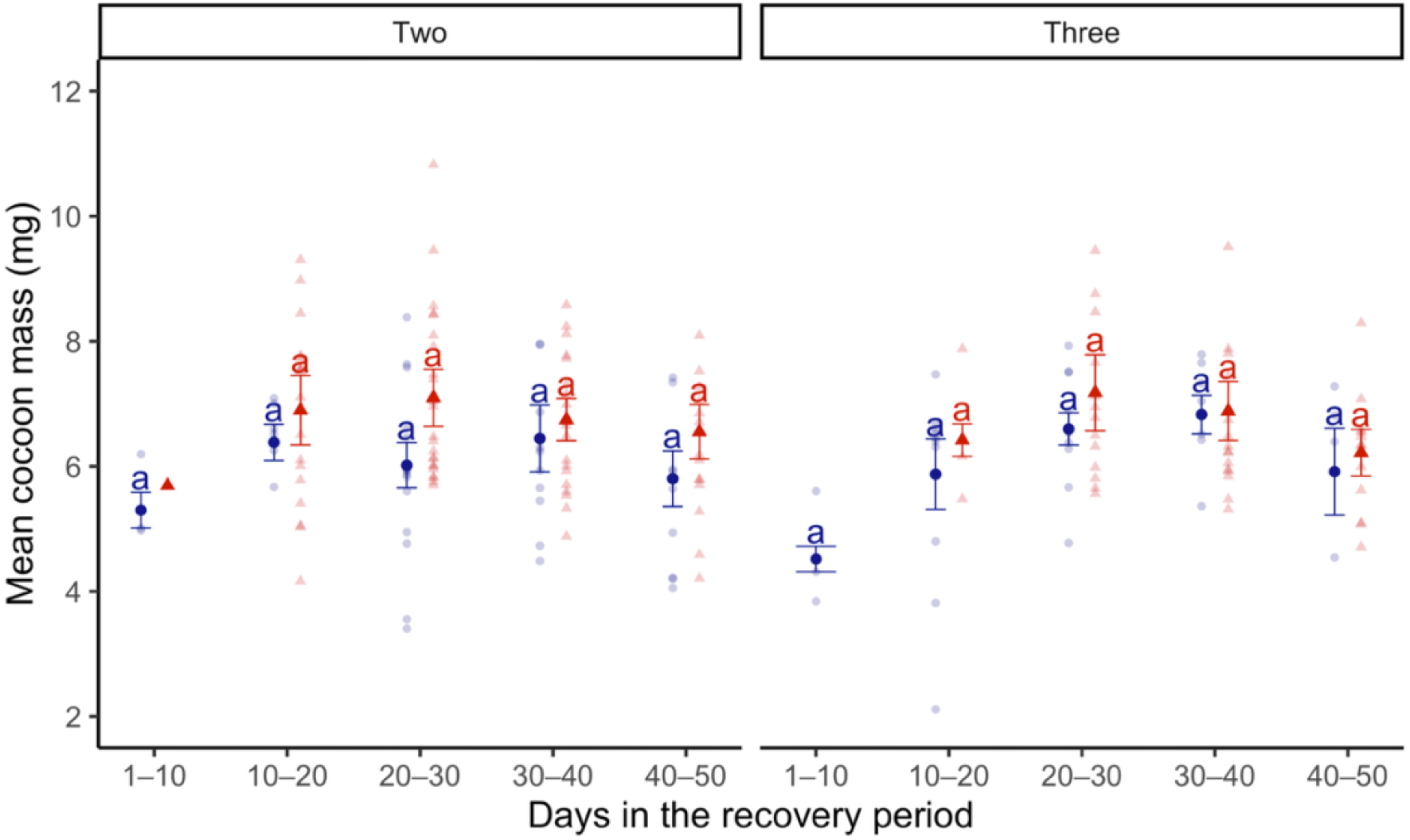
The individual (pale points) and mean (bold points) mass of cocoons produced per group for each of the five 10-day intervals in the recovery period following exposure to constant optimal (blue circles, n = 4) or drought (red triangles, n = 5) conditions, grouped by number of 14-day bouts of drying (or control) conditions experienced. Cocoon mass was not measured for earthworms that experienced a single bout of drying or control conditions. Error bars show standard errors. Treatments with the same letter do not differ to a statistically significant degree (p > 0.05).

### Relationship between earthworm mass and reproductive output

The number of drying bouts (or constant conditions) significantly influenced the mass of clitellate *Al. chlorotica* during the recovery period, with individuals exposed to a single bout of drying (or constant) conditions being significantly heavier than those subjected to two or three bouts (ANOVA: F = 11.53, df = 2, 133, p < 0.001). Clitellate earthworm mass also varied with recovery duration (ANOVA: F = 8.266, df = 1, 133, p < 0.01), increasing from Day 10 to Day 20, peaking at Days 20-30, and declining over days 30 to 50 (**Fig. 7**). Only the three-bout group showed a significant decline from Day 10 to Day 50 (Tukey, p < 0.05). Overall, drought-exposed earthworms were significantly heavier than those maintained under constant control conditions throughout (ANOVA: F = 65.363, df = 1, 133, p < 0.001). These differences were smallest in individuals that had undergone only a single 14-day bout. Notably, after 10 days in the recovery period, the mean mass of *Al. chlorotica* subjected to three drying bouts was significantly higher than that of earthworms from the constant control conditions (Tukey, p < 0.05).

**Figure 7.**
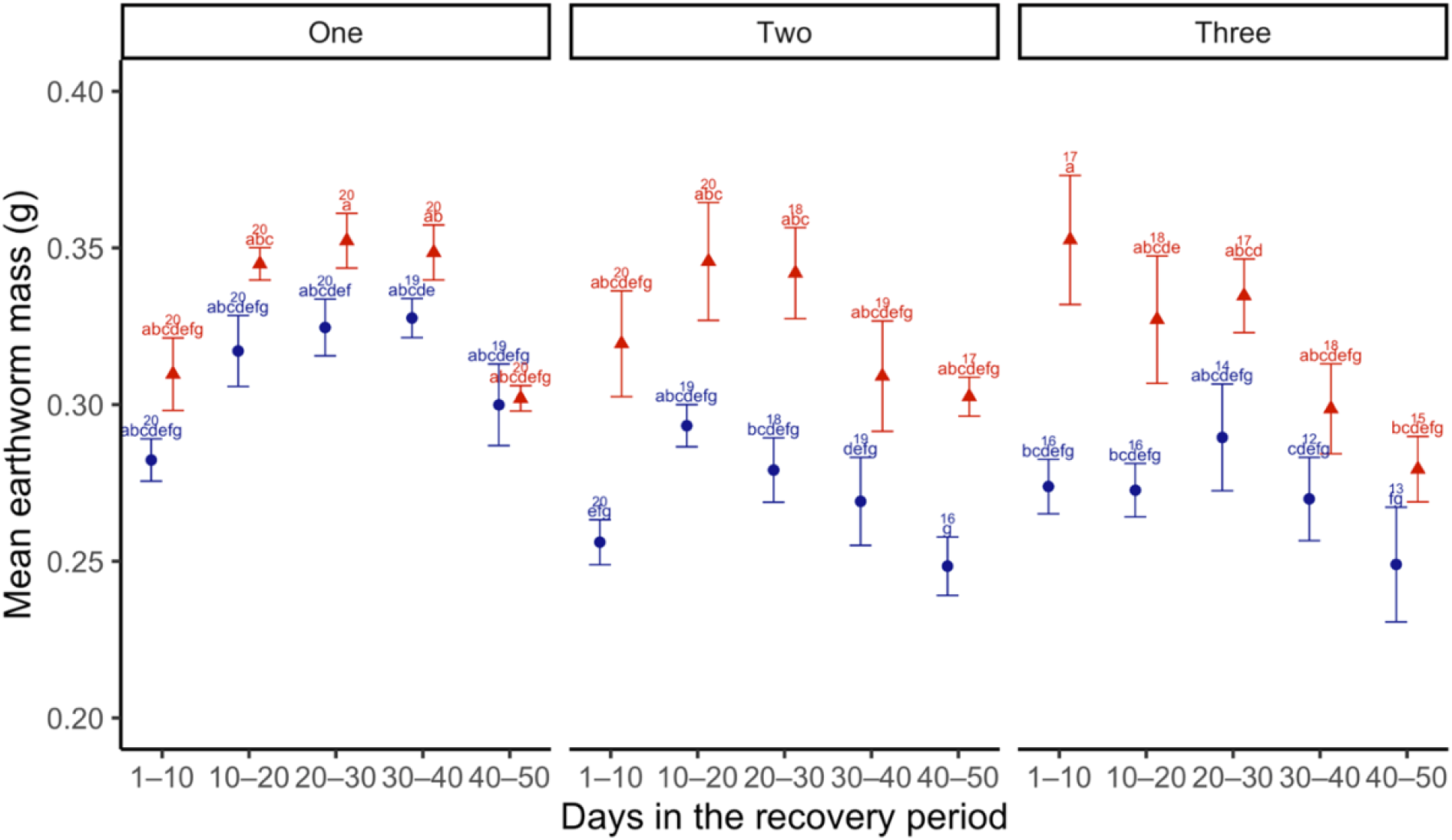
The mean mass of clitellate earthworms (per group) at each of the 10-day sampling points during the recovery period for earthworms exposed to one, two, or three bouts of drying (red triangles, n = 5) or constant moisture conditions (blue circles, n = 4). Treatments with different letters differ to a statistically significant degree (p < 0.05). Numbers represent live clitellate earthworms.

Clitellate earthworm mass correlated positively with cocoon number in both controls (Spearman’s rho = 0.699, n = 76, p < 0.01) and drought treatments (Spearman’s rho = 0.225, n = 76, p = 0.052) (**Fig. 8**). Clitellate earthworm mass was also positively associated with cocoon mass in both drought (Spearman’s rho = 0.592, n = 76, p < 0.01) and control groups (Spearman’s rho = 0.331, n = 76, p < 0.05) (**Fig. 9**).

**Figure 8.**
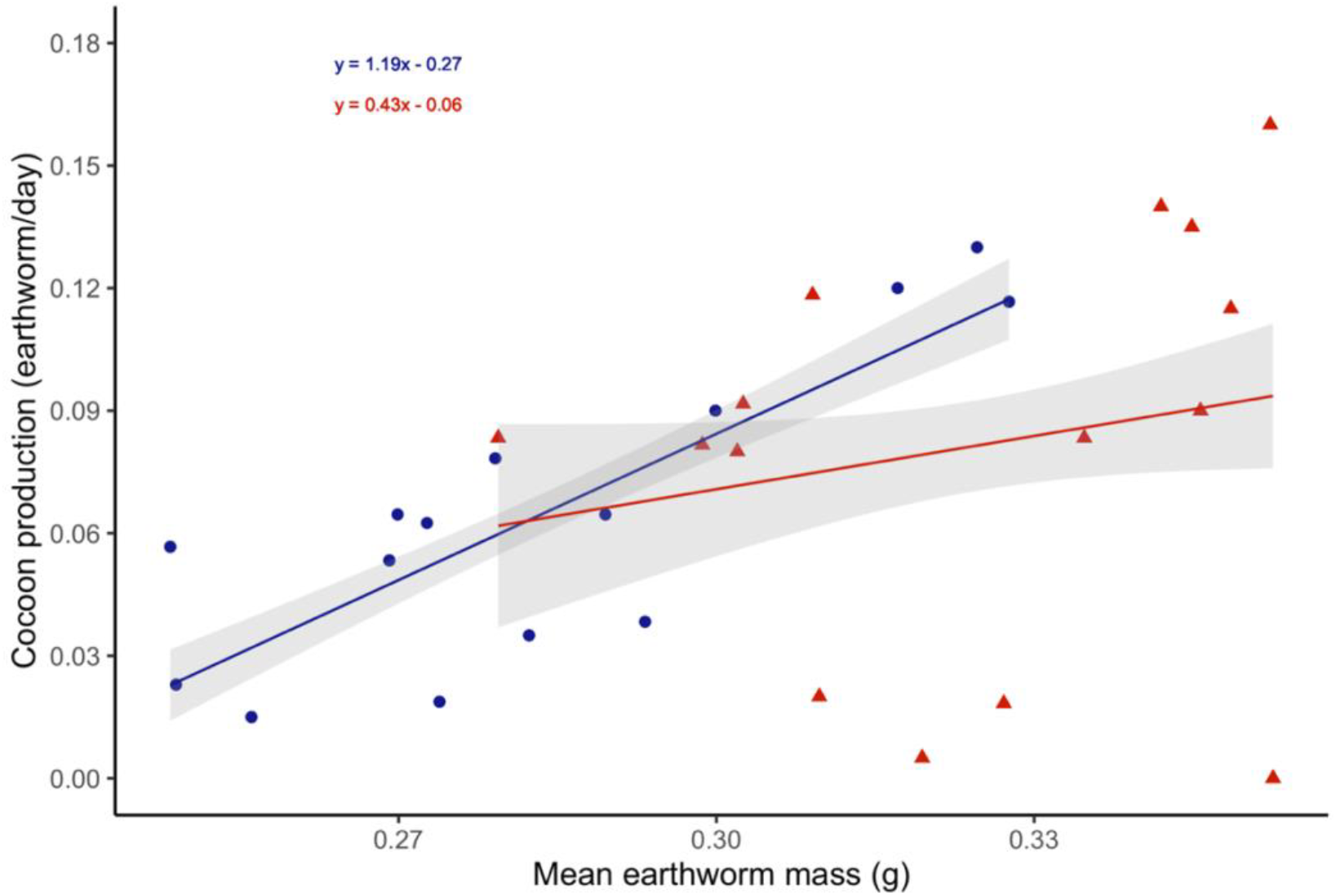
The relationship between the mean mass of clitellate earthworms (per group) and the number of cocoons produced (earthworm/day) in the recovery period following exposure to constant optimal (blue circles) or drying (red triangles) conditions. Lines are fitted using the linear model method with standard error ribbons and equations given for each treatment.

**Figure 9.**
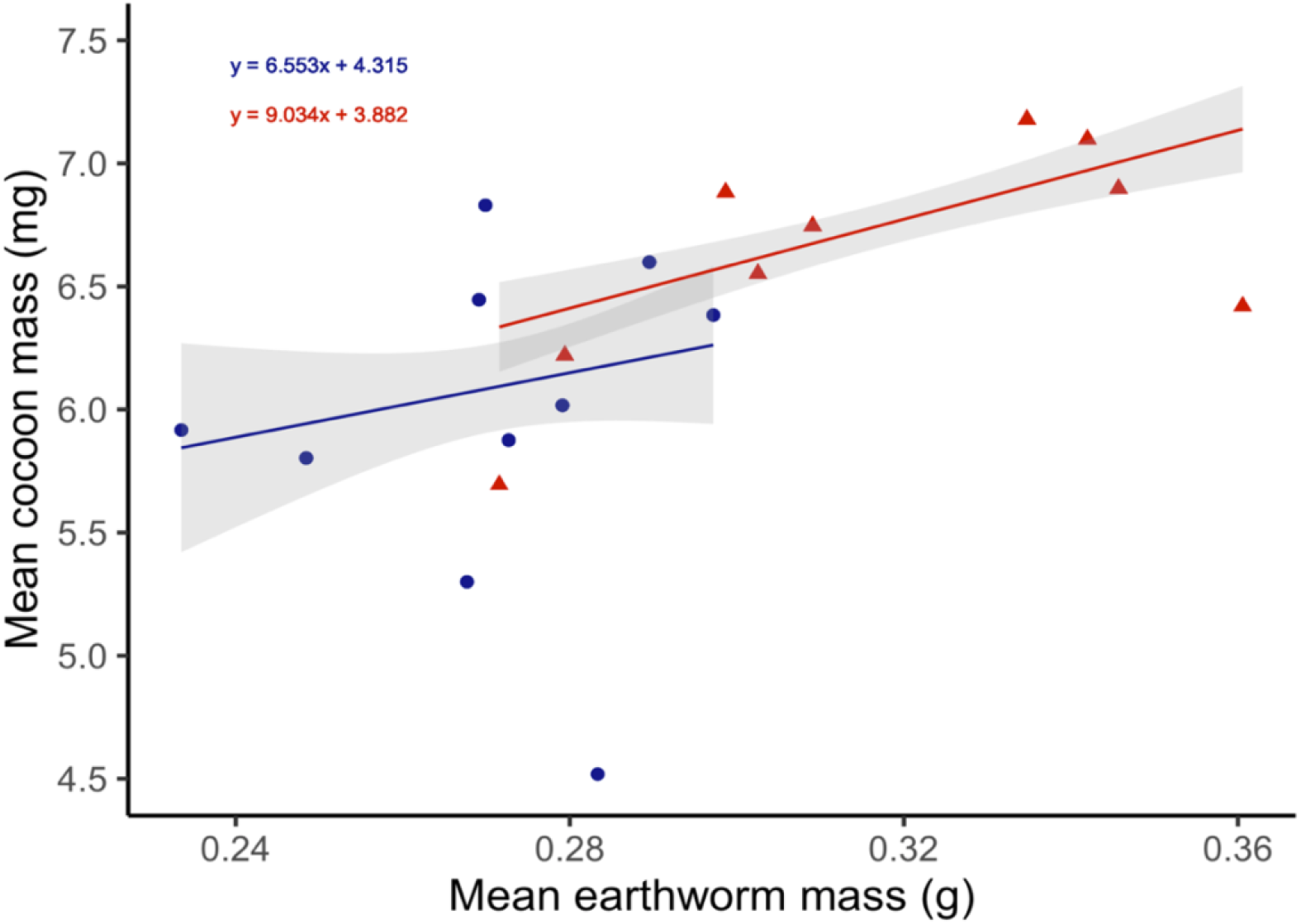
The relationship between the mean mass of clitellate earthworms (g) and the mean mass of cocoons (mg) they produced in the recovery period following exposure to constant optimal (blue circles) or drying (red triangles) conditions. Lines are fitted using the linear model method with standard error ribbons and equations given for each treatment.

### Cocoon viability and development time

Single hatchlings emerged from individual cocoons, consistent with previous findings (Evans and Guild, 1948; Butt, 1997; Butt *et al*., 1997). Both hatching success and incubation time were significantly affected by the number of drying or constant control bouts experienced by the parent earthworms. Overall, *Al. chlorotica* that had undergone three bouts produced cocoons with a higher hatching success (ANOVA: F = 4.897, df = 1, 57, p < 0.05) and shorter incubation time (ANOVA: F = 57/733, df = 1, 75, p < 0.001) compared to those produced by earthworms subjected to two bouts. The timing of cocoon production within the recovery period also influenced outcomes. Both hatching success (ANOVA: F = 5.81, df = 4, 57, p < 0.01) and incubation time (ANOVA: F = 13.409, df = 1, 75, p < 0.001) varied significantly with recovery duration. Cocoons produced during the first 20 days had relatively low viability and longer development times, whereas the highest hatching success (**Fig. 10**) and fastest hatching (**Fig. 11**) occurred in cocoons produced on Days 30-40 (two bouts) and Days 20-40 (three bouts). No significant differences were detected in hatching success (ANOVA: F = 0.08, df = 1, 57, p = 0.778) or incubation time (ANOVA: F = 2.749, df = 1, 75, p = 0.102) between drought-exposed and control treatments overall. Cocoon mass was positively correlated with hatching success for both drought-exposed (Linear regression: F = 15.669, df = 1, 36, p < 0.001) and control groups (Linear regression: F = 12.156, df = 1, 36, p < 0.01) (**Fig. 12**).

**Figure 10.**
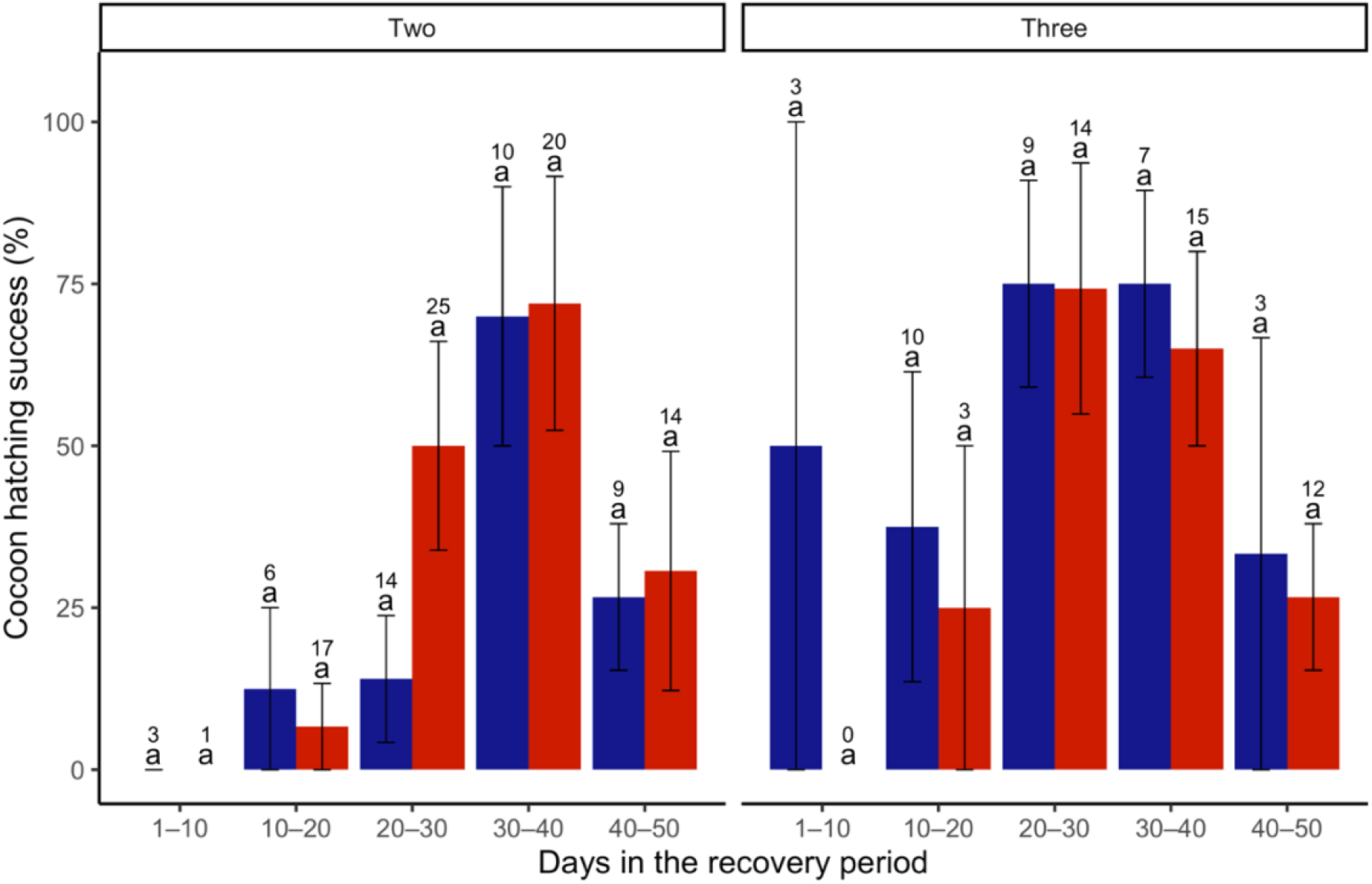
The mean hatching success (percentage hatched) of cocoons produced by replicate groups in each of the 10-day intervals of the recovery period from earthworms previously subjected to either two or three 14-day bouts of constant control (blue) or drying (red) moisture conditions. Error bars show standard error, numbers above bars indicate total numbers of cocoons produced (hatched and unhatched). Treatments with the same letter do not differ to a statistically significant degree (Tukey, p > 0.05).

**Figure 11.**
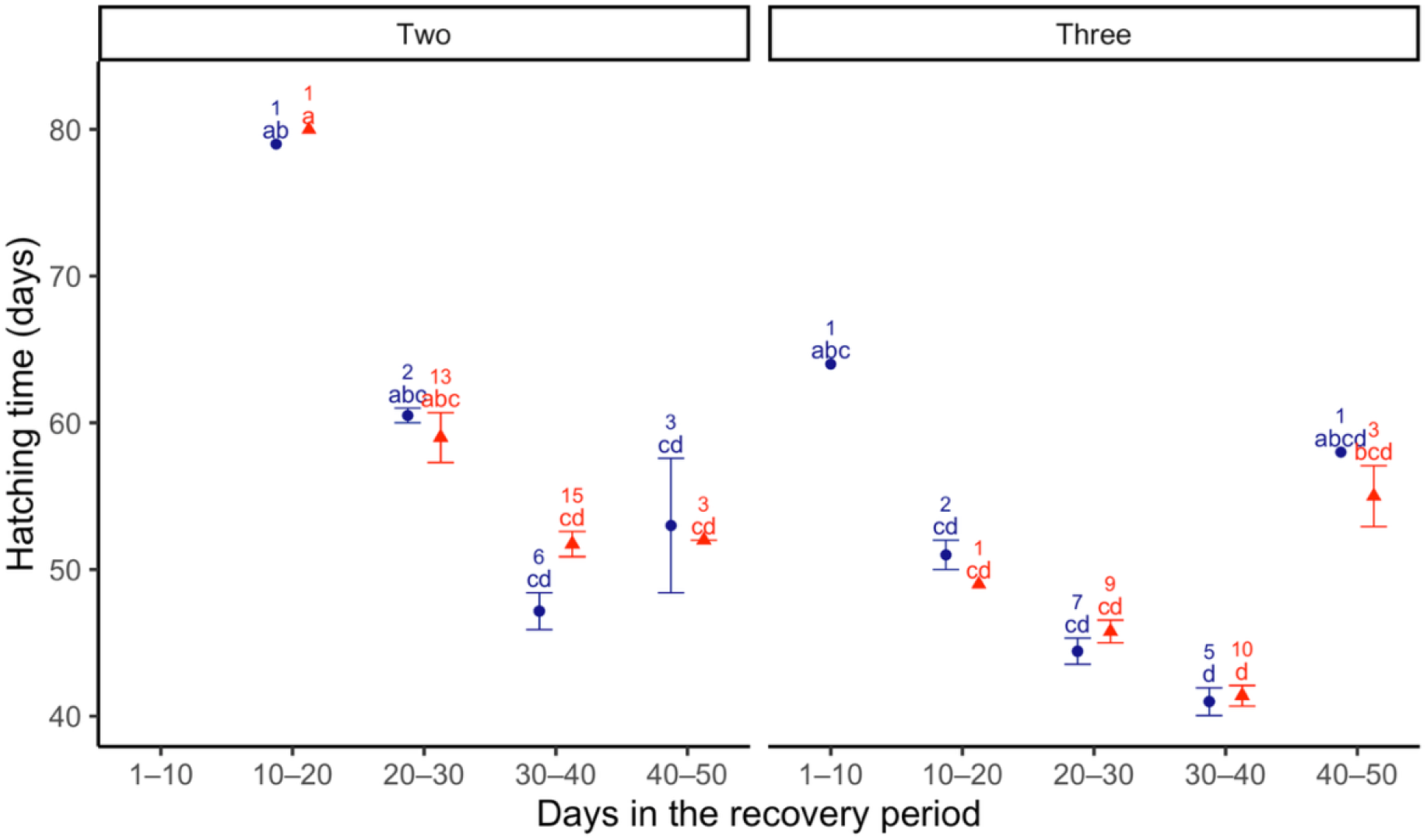
The mean time (days since removal from the experiment) taken for hatchlings to emerge from cocoons produced at each of the 10-day intervals during the recovery period, grouped by those subjected to two or three bouts of constant control (blue circles) or drying (red triangles) conditions. Error bars represent standard error. Treatments with different letters differ to a statistically significant degree (Tukey, p < 0.05). Numbers represent hatched cocoons.

**Figure 12.**
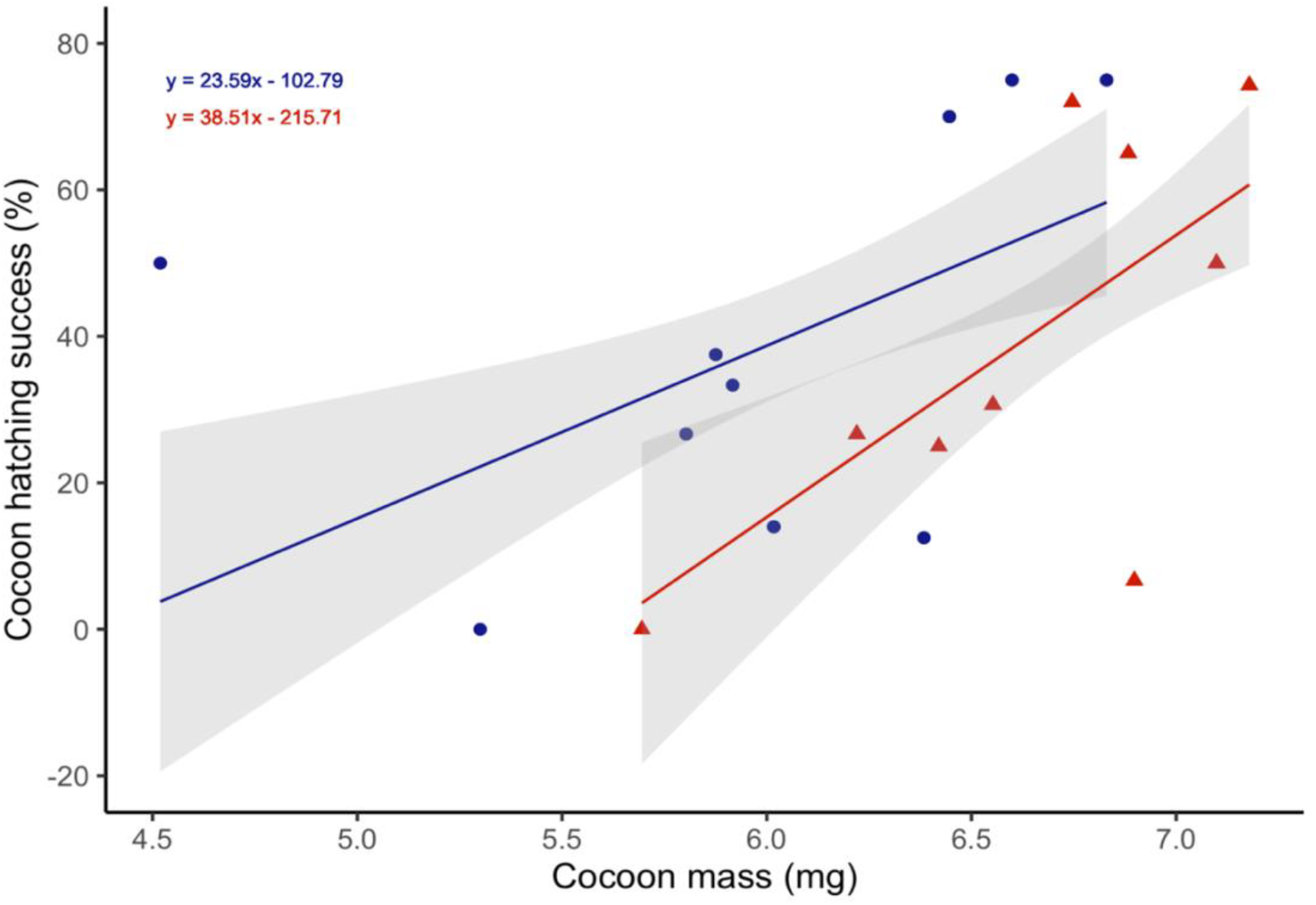
The relationship between the mean mass and the mean hatching success (%) of cocoons produced by replicate groups in the recovery period following exposure to constant optimal (blue circle) or drying (red triangle) conditions. Lines are fitted using the linear model method with standard error ribbons, and equations given for each treatment.

## Discussion

As predicted, repeated drying and associated aestivation significantly reduced the reproductive output of *Al. chlorotica*. Earthworms that experienced a single drying event and thus one period of aestivation, were significantly heavier and produced significantly more cocoons than those exposed to two or three bouts, indicating that *Al. chlorotica* reduce their reproductive investment under repeated stress. Comparable reductions have been observed in other earthworms, for example, *Eisenia fetida* exposed to mechanical stress (experimentally induced contractile movements) produced ∼25 % fewer and ∼30 % lighter cocoons (Aira *et al*., 2007).

Consistent with expectations, the initial rate of cocoon production was low, with fewer and lighter cocoons produced during Days 1-10 of the recovery period. Production increased in Days 10-20 and 20-30 for individuals subjected to one or two bouts of aestivation, respectively. No cocoons were produced during Days 1-10 by earthworms that had experienced three bouts of drying. This delayed recovery parallels the findings of Holmstrup (2001), who reported that *Aporrectodea caliginosa* required 2-4 weeks to resume normal reproduction after exposure to ∼13.2 ± 0.2 wt% (∼12 kPa) to ∼11.1 ± 0.05 wt% (∼22 kPa), despite the earthworms recovering their fresh weight in this time (Holmstrup, 2001). Under more extreme desiccation (∼6.9 ± 0.2 wt%, ∼330 kPa), cocoon production was still reduced even after two months in optimal conditions (Holmstrup, 2001). In contrast, *Al. chlorotica* in the present study resumed cocoon production more rapidly, likely due to milder and more gradual drying employed. Species-specific sensitivity to soil moisture and desiccation may also contribute to the observed differences. *Al. chlorotica* can survive body water losses up to 75 %, compared to 70 % *Lumbricus terrestris* (Roots, 1956). Moreover, the response of earthworms to drying conditions is also likely in part dependent on the conditions they are locally adapted to. Considering this, Holmstrup (2001) acknowledged that the abrupt water potential shifts imposed on *Ap. caliginosa* likely amplified desiccation stress relative to gradual field conditions. The rate of water loss is also critical for cocoon survival, with Petersen *et al*. (2008) suggesting that studies using acute drought exposure may underestimate the tolerance of cocoons to drought conditions. Similar dynamics likely apply to adults, emphasising the ecological relevance of simulating gradual drying regimes. In the present study, soil moisture was reduced gradually from 25 wt% (acclimation) to 20 wt% and then declined progressively to ∼11-13 %. However, *Al. chlorotica* aestivation was abruptly terminated by manual removal, whereas in nature earthworms would resume activity only after conditions became more optimal i.e. when they absorb water from the environment (Jiang *et al*., 2023). A more gradual emergence from aestivation may allow reactivation of metabolism and restore reproductive capacity more smoothly.

Despite differences in water availability, patterns of reproductive output were broadly similar for *Al. chlorotica* from both the drying and constant moisture conditions. Earthworms in the control treatment also showed reduced cocoon production after repeated handling cycles, and no significant differences were observed in cocoon number, mass, or hatching success between treatments. The initially low reproductive output likely reflected limited mating opportunities early in recovery rather than direct effects of prior desiccation. Multiple mating events are known to enhance reproductive success as *Al. chlorotica* produced more juveniles after multiple matings (Dupont *et al*., 2022) and *Eisenia andrei* cocoon viability increased following sperm exchange with several partners (Porto *et al*., 2012). The later decline in cocoon production in the final 10 days of the recovery period may reflect cumulative energetic costs and a reduced proportion of clitellate individuals as some individuals had regressed their sexual characters.

Unexpectedly, *Al. chlorotica* that experienced drying were significantly heavier and produced more, heavier cocoons than earthworms maintained under constant control moisture overall, although cocoon viability was similar across treatments. Comparable compensatory growth responses have been observed by Aira *et al*. (2007) who found that juvenile *E. fetida* exposed to stress subsequently grew faster than controls, while their cocoons had a similar viability, suggesting no effect of stress on cocoon quality. *Al. chlorotica* may have exhibited compensatory feeding once rehydrated as ingestion of food is easier at higher moisture levels (Holmstrup, 2001). Although individuals lost mass during each 14-day drying phase, they regained and exceeded their starting mass after subsequent rewetting, indicating resilience to transient desiccation.

Alternatively, the control conditions may have been suboptimal for growth and reproduction. The mean masses of cocoons produced by *Al. chlorotica* in the drying and control treatments were 6.701 ± 1.205 mg (n = 74) and 6.036 ± 1.256 mg (n = 121) respectively. These are lower than published averages for *Al. chlorotica* cocoons of 7 mg (in experimental conditions with a 8:5 ratio of soil to separated cattle solids at 10-20 °C, soil moisture content not reported, Butt, 1997), ∼9 to 16 mg (in experiments with field soils kept at ∼25 and 43 % moisture respectively, exposed variations in temperature from 10 to 8 °C, Evans and Guild, 1948) and 11 ± 1 mg (for earthworms kept at 15 °C in soil with moist cow dung, soil moisture content not specified, Holmstrup, 1994). Differences in soil conditions including the source and quantity of supplemented organic matter, could contribute to the varying earthworm masses. Moreover, overall hatching success was relatively low in both treatments, at 45.5 % (55/121) for the drying treatment and 37.8 % (28/74) for the control group. Previous studies report highly variable *Al. chlorotica* hatching success ranging from 34 % to ∼90 % depending on rearing conditions (Pedersen and Bjerne, 1991; Jensen and Holmstrup, 1997; Butt, 1997; Butt *et al*., 1997). The observation that four control individuals entered aestivation suggests that despite adequate moisture (∼25 wt%), the conditions were suboptimal. Potvin and Lilleskov (2017) noted that earthworms may aestivate to avoid starvation in the absence of appropriate food availability, therefore, limited organic matter could be a contributing factor. Consistent with this, control earthworms began to lose mass after the first 14-day period and eventually fell below their starting mass, indicating energy limitation. When other physiochemical environmental factors are suitable, food quality and quantity are the main constraints on earthworm populations (Curry, 2004). Similarly, Diehl and Williams (1992) demonstrated that growth and activity of *E. fetida* depend on both soil moisture and food availability.

In this experiment, organic matter was incorporated prior to earthworm introduction and not replenished during the 14-day periods. Thus, the control soils likely experienced continuous food depletion as it is easier for earthworms to ingest food in higher moisture contents (Holmstrup, 2001), whereas individuals under drying conditions would have ceased feeding when they entered aestivation (Díaz Cosín *et al*., 2006). Additionally, the higher soil moisture in the control conditions and recovery period may have also created more hospitable conditions for soil microbes (Cook and Orchard, 2008; Liu *et al*., 2022), accelerating organic matter decomposition and potentially leaving food of lower quality or reduced digestibility for earthworms. However, if food availability had been the limiting factor, an increase in earthworm mass might have been expected following transfers to fresh soils where food was more abundant i.e. in the 3-day transitions between each 14-day period and every 10 days in the recovery period. This was not observed, instead changes in mass progressively declined with increasing time spent in the recovery period, even for earthworms previously exposed to drying (**Fig. S3**). In addition, organic matter remained visible in soils at the end of each exposure period, suggesting that food remained available, although it may have been reduced quality or indigestible.

The gravimetric soil moisture of 25 wt% selected for the control treatment followed recommended culture conditions (Lowe and Butt, 2005). However, *Al. chlorotica* green morph may prefer and perform better at higher moisture levels. Lowe and Butt (2007) reported significantly higher growth and maturation rates in hatchlings at 29 wt% compared with 21 wt% moisture. Similarly, Satchell (1967) found green morphs dominated (>90 %) in sites >40 % moisture, while pink morphs prevailed below 25 %. Thus, 25 wt% moisture likely represents the lower tolerance limit for the green morph, sufficient for activity but not optimal for reproduction. Moreover, although drought-exposed earthworms were exposed to more severe water limitation, they may have largely avoided its costs by entering aestivation, in which gas exchange is depressed to conserve energy and body fluid osmolality increases to prevent water loss (Bayley *et al*., 2010). Upon reactivation, they may have had greater residual fitness than controls that remained active under suboptimal conditions. In addition, repeated DIW surface additions caused compaction of the control soils, likely due to the breakdown of weaker soil aggregates. Increased bulk density reduces pore space and raises mechanical resistance, constraining burrowing and increasing energy costs in earthworms such as *Ap. longa* and *Ap. caliginosa* (Arrázola-Vásquez *et al*., 2022). Structural changes in the control soils resulting in reduced pore spaces may therefore have further contributed to their suboptimal nature.

Earthworm mass strongly influenced reproductive output. Cocoon number, cocoon mass, and hatching success all peaked between Days 20-40 of the recovery period when earthworms were heaviest. Positive relationships between adult size and cocoon output are well documented. For instance, Teferedegn and Ayele (2024), found cocoon production in three epigeic species increased initially but declined after ∼13 to ∼17 days, coinciding with progressive weight loss of the earthworms. Similar positive associations between earthworm mass and cocoon production have been reported for *E. andrei* (Van Gestel *et al*., 1992) and for earthworm and cocoon mass in eight tropical earthworm species (Chaudhuri and Bhattacharjee, 2011). Holmstrup (2001) proposed that water loss cues reproductive inhibition, as cocoon production declines at water potentials when earthworms remain active but lose body mass. In contrast to our findings, Pedersen and Bjerre (1991) found no relationship between cocoon mass and hatching success in *Al. chlorotica, Ap. longa, Ap. caliginosa* or *L. terrestris* cocoons, although they did find a significant positive correlation between cocoon and hatchling mass.

In the present study, the mean incubation time (50.61 ± 8.45 days) was consistent with previous reports of 50 (Gerard, 1967), 59 (Holmstrup *et al*., 1991) and 51.4 (Butt, 1997) days at the same temperature (15 °C). However, there was a significant effect of time in the recovery period as cocoons produced in the first 20 days of the recovery period were slower to hatch than those produced on Days 30-40. This along with their smaller mass and lower viability suggests reduced reproductive quality immediately following exposure to stressful conditions. This suggests that at least ∼50 days of optimal moisture may be required for successful *Al. chlorotica* cocoon development following drought, although this will likely depend on the soil, namely temperature. For instance, Holmstrup *et al*. (1991) found that *Al. chlorotica* cocoons took 34-38 days to develop at 20 °C, while at 5 °C incubation exceeded 400 days. Further work could examine whether aestivation negatively affects hatchling growth to better understand the duration of favourable conditions required for earthworm populations to recover from exposure to stressful conditions.

Regression and reabsorption of secondary sexual structures have been reported to be a key feature of aestivation (Olive and Clark, 1978; Jiménez *et al*., 2000). These changes in sexual mating characteristics resulting from exposure to stressful conditions are temporary and functions are usually regained on return to favourable conditions (Christyraj *et al*., 2025). This allows allocation of resources for survival to be prioritised over current reproduction in the hopes of survival and reproduction in future improved conditions. Evidence of such a trade-off was observed in this experiment, as some adult *Al. chlorotica* regressed their clitellum, usually associated with individuals below 0.2 g in mass. This occurred not only in drying treatments but also in active earthworms in the constant higher moisture conditions, particularly during the 50-day recovery period, again suggesting that control conditions were suboptimal. Butt (1997) similarly found that the reproductive state of *Al. chlorotica* was strongly influenced by environmental conditions, with only 23.3 % of the adults remaining clitellate after one year at 10 °C, compared to 60 % at 15 °C and 86.7 % at 20 °C. While reproductive output in the present study was adjusted to account for the reduction of clitellate individuals, some individuals regained their fully developed state during the recovery period, demonstrating plasticity. Future research is needed to investigate whether regression and re-development of the clitellum impose lasting costs on reproductive success.

## Conclusion

Consistent with previous studies, *Al. chlorotica* exposed to multiple bouts of drying and aestivation produced fewer and lighter cocoons than those exposed to a single bout, indicating a reduction in reproductive output under repeated stress. However, contrary to expectations, earthworms from drying treatments generally recovered mass more effectively and produced more and heavier cocoons than those maintained under constant control conditions. This suggests that aestivation confers an adaptive benefit as a protective strategy and that subsequent recovery under favourable conditions can stimulate compensatory growth and reproduction. The lower mass and reproductive output of earthworms under constant moisture conditions indicate that, while not water-limited, control conditions were suboptimal, likely due to progressive depletion of organic matter and increased soil compaction constraining feeding and movement. This emphasises the importance of considering soil quality and structure alongside moisture availability when evaluating drought effects on earthworms. Across treatments, earthworm body mass strongly predicted fecundity, with maximum cocoon production, cocoon mass, and hatching success occurring when earthworms were heaviest. Nevertheless, cocoon quality and incubation time varied over the recovery period, with reduced performance in the first 20 days suggesting a lag in reproductive recovery following stress. Evidence of clitellum regression further indicates that both drying and constant-moisture conditions imposed physiological stress that temporarily shifted resource allocation away from reproduction. Overall, these findings demonstrate that *Al. chlorotica* are resilient to short-term, intermittent drought through aestivation, but that reproductive output is sensitive to the combined effects of soil moisture, food availability and soil physical structure. Increasing drought frequency and duration are therefore likely to intensify negative impacts on population dynamics. For populations to persist under fluctuating environmental conditions, sustained periods of favourable conditions are essential to enable recovery of both adult condition and cocoon viability. Further research could investigate the long-term reproductive consequences of aestivation, as well as the vulnerability of earthworm species lacking this adaptive strategy.

## Acknowledgements

The study was supported by an Adapting to the Challenges of a Changing Environment Doctoral Training Partnership studentship (NE/S00713X/1).

## Author contributions

Conceptualisation, RVAB, PJW, MEH; Investigation, RVAB; Methodology, RVAB, PJW, MEH; Supervision, PJW, MEH; Formal analysis, RVAB; Writing – original draft, RVAB; Writing – review and editing, RVAB, PJW, MEH.

## References

Agrigem. Lincoln LN1 2FU. Available at: https://www.agrigem.co.uk/product/kettering-loam-25kg/?gad_source=1&gbraid=0AAAAADuriRyu7EkhqjUR1clb63YjKeSbo&gclid=Cj0KCQjw_JzABhC2ARIsAPe3ynqNUsp-t-kjVYmzBZfuGgYw4EwcdB7jBlik34uQsgKWDw9klQD6WtAaAoUBEALw_wcB

Aira, M., Domínguez, J., Monroy, F., and Velando, A. (2007). Stress promotes changes in resource allocation to growth and reproduction in a simultaneous hermaphrodite with intermediate growth. Biological Journal of the Linnean Society. 91(4), 593–600. 10.1111/j.1095-8312.2007.00822.x

Arrázola-Vásquez, E., Larsbo, M., Capowiez, Y., Taylor, A., Sandin, M., Iseskog, D., and Keller, T. (2022). Earthworm burrowing modes and rates depend on earthworm species and soil mechanical resistance. Applied Soil Ecology : A Section of Agriculture, Ecosystems and Environment. 178. 10.1016/j.apsoil.2022.104568

Bart, S., Amossé, J., Lowe, C. N., Mougin, C., Péry, A. R. R., and Pelosi, C. (2018). Aporrectodea caliginosa, a relevant earthworm species for a posteriori pesticide risk assessment: current knowledge and recommendations for culture and experimental design. Environmental Science and Pollution Research International. 25(34), 33867–33881. 10.1007/s11356-018-2579-9

Bayley, M., Overgaard, J., Høj, A. S., Malmendal, A., Nielsen, N. C., Holmstrup, M., and Wang, T. (2010). Metabolic Changes during Estivation in the Common Earthworm Aporrectodea caliginosa. Physiological and Biochemical Zoology. 83(3), 541–550. 10.1086/651459

Blouin, M., Hodson, M. E., Delgado, E. A., Baker, G., Brussaard, L., Butt, K. R., Dai, J., Dendooven, L., Peres, G., Tondoh, J. E., Cluzeau, D., and Brun, J.-J. (2013). A review of earthworm impact on soil function and ecosystem services. European Journal of Soil Science. 64, 161–182. 10.1111/ejss.12025

Butt, K. R. (1997). Reproduction and growth of the earthworm Allolobophora chlorotica (Savingy, 1826) in controlled environments. Pedobiologia, 41(4), 369– 374. 10.1016/S0031-4056(24)00253-1

Butt, K. R., Frederickson, J., and Morris, R. M. (1997). The Earthworm Inoculation Unit technique: An integrated system for cultivation and soil-inoculation of earthworms. Soil Biology and Biochemistry. 29(3), 251–257. 10.1016/S0038-0717(96)00053-3

Chaudhuri, P. S., and Bhattacharjee, S. (2011). Reproductive biology of eight tropical earthworm species of rubber plantations in Tripura, India. Tropical Ecology. 52(1), 49–60.

Christyraj, S. J. D., Vaidhyalingham, A. B., Sengupta, C., Rajagopalan, K., Vadivelu, K., Suresh, N. K., and Venkatachalam, B. (2025). Functional significance of earthworm clitellum in regulating the various biological aspects of cell survival and regeneration. Developmental Dynamics. 254(3), 212–221. 10.1002/dvdy.751

Cook, F. J., and Orchard, V. A. (2008). Relationships between soil respiration and soil moisture. Soil Biology and Biochemistry. 40(5), 1013–1018. 10.1016/j.soilbio.2007.12.012

Curry, J. P. (2004). Factors affecting the abundance of earthworms in soils. In Earthworm Ecology, Second Edition (pp. 91–113). 10.1201/9781420039719

Dai, A. (2013). Increasing drought under global warming in observations and models. Nature Climate Change. 3(2), 171. 10.1038/nclimate1811

Diehl, W. J., and Williams, D. L. (1992). Interactive effects of soil moisture and food on growth and aerobic metabolism in eisenia fetida (oligochaeta). Comparative Biochemistry and Physiology. A, Comparative Physiology. 102(1), 179–184. 10.1016/0300-9629(92)90031-K

Dupont, L., Audusseau, H., Porco, D., and Butt, K. R. (2022). Mitonuclear discordance and patterns of reproductive isolation in a complex of simultaneously hermaphroditic species, the Allolobophora chlorotica case study. Journal of Evolutionary Biology. 35(6), 831–843. 10.1111/jeb.14017

Edwards, C. A. and Bohlen, P. J. Biology and Ecology of Earthworms. 3rd ed. / C. A. Edwards and P.J. Bohlen. London: Chapman and Hall, (1996). Print.

Evans, A. C., and Guild, W. J. M. (1948). Studies on the relationships between earthworms and soil fertility. Annals of Applied Biology. 35(4), 471– 484. 10.1111/j.1744-7348.1948.tb07391.x

Fründ, H-C., Butt, K., Capowiez, Yvan., Eisenhauer, N., Emmerling, C., Ernst, Gregor., Potthoff, M., Schädler, M., Schrader, S. (2010). Using earthworms as model organisms in the laboratory: Recommendations for experimental implementations. Pedobiologia. 53(2), 119–125. 10.1016/j.pedobi.2009.07.002

Gerard, B. M. (1967). Factors affecting earthworms in pastures. Journal of Animal Ecology. 36, 235–252. 10.2307/3024

Holden, J., Grayson, R. P., Berdeni, D., Bird, S., Chapman, P. J., Edmondson, J. L., Firbank, L.G., Helgason, T., Hodson, M. E., Hunt, S.F.P., Jones, D.T., Lappage, M.G., Marshall-Harries, E., Nelson, M., Prendergast-Miller, M., Shaw, H., Wade, R.N and Leake, J. R. (2019). The role of hedgerows in soil functioning within agricultural landscapes. Agriculture, Ecosystems and Environment. 273, 1–12. 10.1016/j.agee.2018.11.027

Holmstrup, M., Østergaard, I.K., Nielsin, A., Hansen, B.T. (1991). The relationship between temperature and cocoon incubation time for some lumbricid earthworm species. Pedobiologia. 35, 179–184.

Holmstrup, M. (1994). Physiology of cold hardiness in cocoons of five earthworm taxa (Lumbricidae: Oligochaeta). Journal of Comparative Physiology. B, Biochemical, Systemic, and Environmental Physiology. 164(3), 222–228. 10.1007/BF00354083

Holmstrup, M. (2001). Sensitivity of life history parameters in the earthworm Aporrectodea caliginosa to small changes in soil water potential. Soil Biology and Biochemistry. 33(9), 1217–1223. 10.1016/S0038-0717(01)00026-8

Jensen, K. S., and Holmstrup, M. (1997). Estimation of earthworm cocoon development time and its use in studies of in situ reproduction rates. Applied Soil Ecology: A Section of Agriculture, Ecosystems and Environment. 7(1), 73–82. 10.1016/S0929-1393(97)00016-4

Jiang, C., Storey, K. B., Yang, H., and Sun, L. (2023). Aestivation in Nature: Physiological Strategies and Evolutionary Adaptations in Hypometabolic States. International Journal of Molecular Sciences. 24(18), 14093. 10.3390/ijms241814093

Jiménez, J. J., Brown, G. G., Decaëns, T., Feijoo, A., and Lavelle, P. (2000). Differences in the timing of diapause and patterns of aestivation in tropical earthworms. Pedobiologia, 44(6), 677–694. 10.1078/S0031-4056(04)70081-5

Kretzschmar, A., and Bruchou, C. (1991). Weight response to the soil water potential of the earthworm Aporrectodea longa. Biology and Fertility of Soils. 12(3), 209–212. 10.1007/BF00337204

Liu, L., Estiarte, M., Bengtson, P., Li, J., Asensio, D., Wallander, H., and Peñuelas, J. (2022). Drought legacies on soil respiration and microbial community in a Mediterranean forest soil under different soil moisture and carbon inputs. Geoderma, 405, 115425. 10.1016/j.geoderma.2021.115425

Lowe, C. N., and Butt, K. R. (2005). Culture techniques for soil dwelling earthworms: A review. Pedobiologia. 49(5), 401–413. 10.1016/j.pedobi.2005.04.005

Lowe, C. N., and Butt, K. R. (2007). Life-cycle traits of the dimorphic earthworm species Allolobophora chlorotica (Savigny, 1826) under controlled laboratory conditions. Biology and Fertility of Soils. 43(4), 495–499. 10.1007/s00374-006-0154-x

McDaniel J, P., Barbarick K, A., Stromberger M, E., and Cranshaw W. (2013a). Survivability of Aporrectodea caliginosa in response to drought stress in a Colorado soil. Soil Science Society of America journal. 77(5), 1667–1672. 10.2136/sssaj2013.02.0064

McDaniel, J. P., M. E. Stromberger, K. A. Barbarick and W. Cranshaw (2013b). Survival of Aporrectodea caliginosa and its effects on nutrient availability in biosolids amended soil. Applied Soil Ecology. 71, 1– 6. 10.1016/j.apsoil.2013.04.010

Nepstad, D. C., Moutinho, P., Dias-Filho, M. B., Davidson, E., Cardinot, G., Markewitz, D., Figueiredo, R., Vianna, N., Chambers, J., Ray, D., Guerreiros, J. B., Lefebvre, P., Sternberg, L., Moreira, M., Barros, L., Ishida, F. Y., Tohlver, I., Belk, E., Kalif, K., Schwalbe, K. (2002). The effects of partial throughfall exclusion on canopy processes, aboveground production, and biogeochemistry of an Amazon forest. Journal of Geophysical Research. D. Atmospheres. 107(D20), LBA 53-1-LBA 53-18. 10.1029/2001JD000360

Olive and Clark (1978). Physiology of reproduction. In Physiology and Annelids (ed. By P. J. Mill), Academic Press, London.

Pedersen, M. B., and Bjerre, A. (1991). The relationship between mass of newly hatched individuals and cocoon mass in lumbricid earthworms. Pedobiologia. 35(1), 35–39. 10.1016/S0031-4056(24)00042-8

Petersen, C. R., Holmstrup, M., Malmendal, A., Bayley, M., and Overgaard, J. (2008). Slow desiccation improves dehydration tolerance and accumulation of compatible osmolytes in earthworm cocoons (Dendrobaena octaedra Savigny). Journal of Experimental Biology. 211(12), 1903–1910. 10.1242/jeb.017558

Porto, P. G., Velando, A., and Domínguez, J. (2012). Multiple mating increases cocoon hatching success in the earthworm Eisenia andrei (Oligochaeta: Lumbricidae): EFFECTS OF MULTIPLE MATING ON E. ANDREI REPRODUCTION. Biological Journal of the Linnean Society. 107(1), 175–181. 10.1111/j.1095-8312.2012.01913.x

Potvin, L. R., and Lilleskov, E. A. (2017). Introduced earthworm species exhibited unique patterns of seasonal activity and vertical distribution, and Lumbricus terrestris burrows remained usable for at least 7 years in hardwood and pine stands. Biology and Fertility of Soils. 53(2), 187–198. 10.1007/s00374-016-1173-x

R Core Team (2024). R: A Language and Environment for Statistical Computing. R Foundation for Statistical Computing, Vienna, Austria. https://www.R-project.org/.

Ramsay, J. A. (1949). The Osmotic Relations of the Earthworm. Journal of Experimental Biology. 26(1), 46–56. 10.1242/jeb.26.1.46

Reinecke, S. A., and Reinecke, A. J. (2007). Biomarker response and biomass change of earthworms exposed to chlorpyrifos in microcosms. Ecotoxicology and Environmental Safety. 66(1), 92–101. 10.1016/j.ecoenv.2005.10.007

Roots, B. I. (1956). The Water Relations of Earthworms. Journal of Experimental Biology. 33(1), 29–44. 10.1242/jeb.33.1.29

Ruiz, S. A., Bickel, S., and Or, D. (2021). Global earthworm distribution and activity windows based on soil hydromechanical constraints. Communications Biology. 4(1), 612–612. 10.1038/s42003-021-02139-5

Satchell J. E. (1967) Lumbricidae. In Soil Biology (A. Burges and F. Raw, Eds), 259–322. Academic Press, London.

Seneviratne, S.I., X. Zhang, M. Adnan, W. Badi, C. Dereczynski, A. Di Luca, S. Ghosh, I. Iskandar, J. Kossin, S. Lewis, F. Otto, I. Pinto, M. Satoh, S.M. Vicente-Serrano, M. Wehner, and B. Zhou, (2021). Weather and Climate Extreme Events in a Changing Climate. In Climate Change 2021: The Physical Science Basis. Contribution of Working Group I to the Sixth Assessment Report of the Intergovernmental Panel on Climate Change [Masson-Delmotte, V., P. Zhai, A. Pirani, S.L. Connors, C. Péan, S. Berger, N. Caud, Y. Chen, L. Goldfarb, M.I. Gomis, M. Huang, K. Leitzell, E. Lonnoy, J.B.R. Matthews, T.K. Maycock, T. Waterfield, O. Yelekçi, R. Yu, and B. Zhou (eds.)]. Cambridge University Press, Cambridge, United Kingdom and New York, NY, USA, pp. 1513–1766, 10.1017/9781009157896.013

Sherlock, E. (2018). Key to the earthworms of the UK and Ireland. Telford: Fsc Publications.

Teferedegn, G. D., and Ayele, C. (2024). Life Cycle Patterns of Epigeic Earthworm Species (Eisenia fetida, Eisenia andrei, and Dendrobaena veneta) in a Blend of Brewery Sludge and Cow Dung. International Journal of Zoology. 1–7. 10.1155/2024/6615245

Tilikj, N., and Novo, M. (2022). How to resist soil desiccation: Transcriptional changes in a Mediterranean earthworm during aestivation. Comparative Biochemistry and Physiology. Part A, Molecular and Integrative Physiology, 264, 111112–111112. 10.1016/j.cbpa.2021.111112

van Gestel, C., Dirven-Van Breemen, E. M., and Baerselman, R. (1992). Influence of environmental conditions on the growth and reproduction of the earthworm Eisenia andrei in an artificial soil substrate. Pedobiologia. 36, 109–120. 10.1016/s0031-4056(24)00779-0

West, H. K., Morgan, A. J., Bowker, D. W., Davies, M. S., and Herbert, R. J. (2003). Evidence for interpopulation differences in life history parameters of adult and F1 generation Lumbricus rubellus. Pedobiologia. 47(5), 535–541. 10.1078/0031-4056-00225

Weil, R. R., and Brady, N. C. (2017). The nature and properties of soils (Fifteenth edition). Harlow, England: Pearson

Whalen, J., Parmelee, R., and Subler, S. (2000). Quantification of nitrogen excretion rates for three lumbricid earthworms using N-15. Biology and Fertility of Soils. 32(4), 347–352. 10.1007/s003740000259

