## Supplementary figures for "The effect of repeated periods of drought and aestivation on *Allolobophora chlorotica* reproductive output"

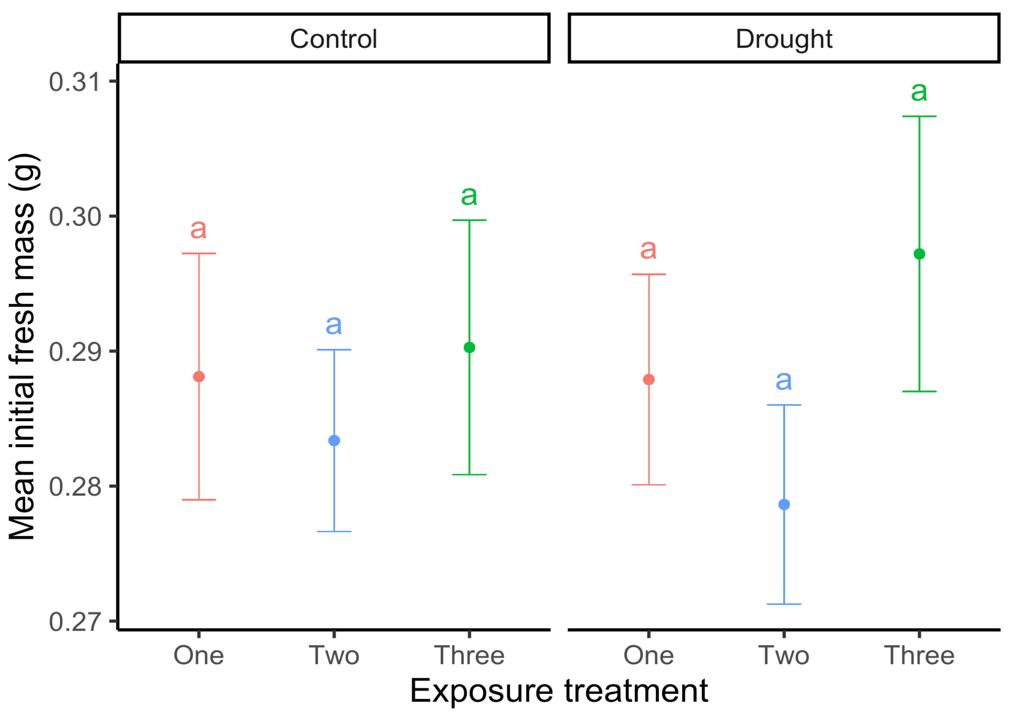

**Figure S1**. The mean (n = 20) initial mass of *Al. chlorotica* assigned to experience either one (red), two (blue) or three (green) bouts of control or drying conditions. Error bars show standard error. Treatments with the same letter do not differ to a statistically significant degree (p > 0.05, Tukey test).

**Figure S2**. The mean overall cocoon production (per earthworm per day) of *Al. chlorotica* assigned to experience either one (red), two (blue) or three (green) bouts of control or drying conditions. Error bars show standard error. Treatments with the same letter do not differ to a statistically significant degree (p > 0.05, Tukey test).

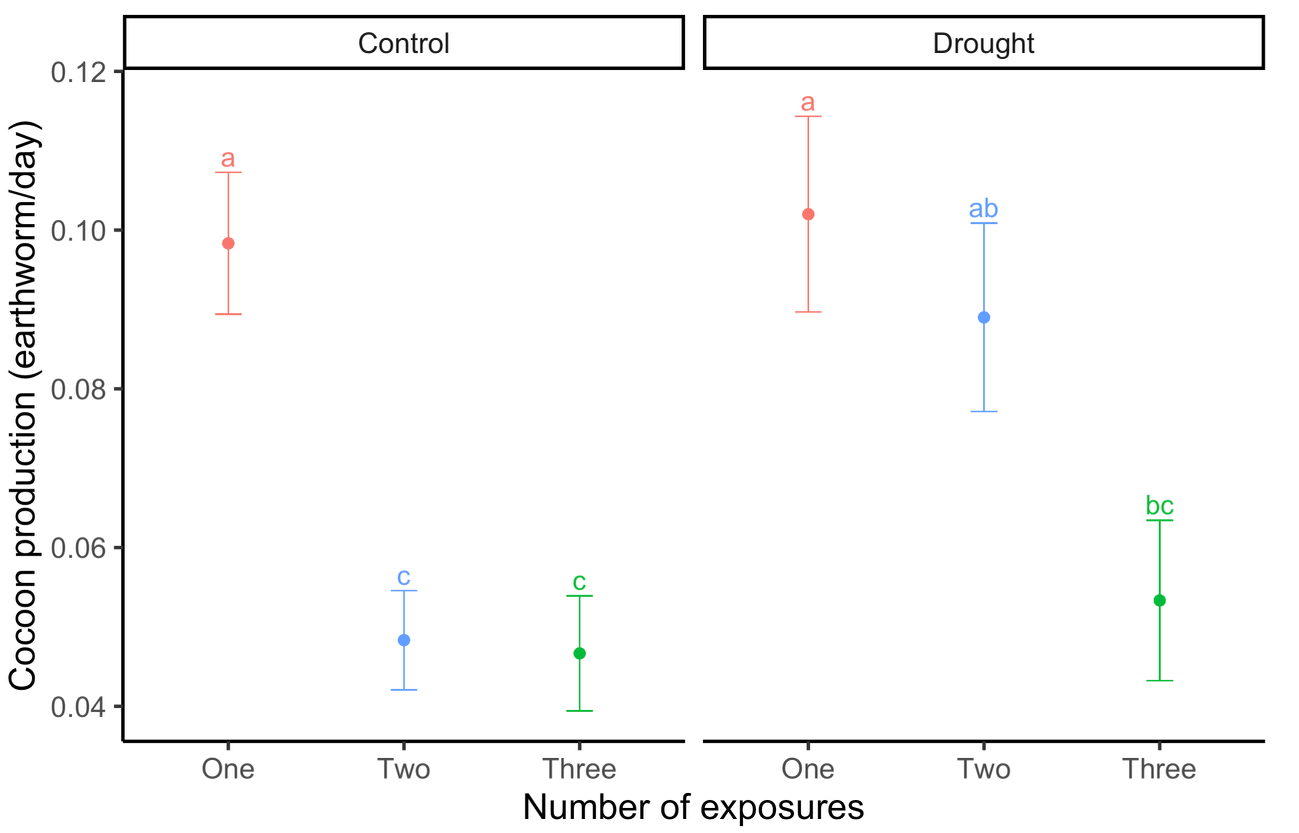

**Figure S3**. The mean change in mass (n = 5) during the recovery period relative to the starting mass of earthworms subjected to one (red), two (blue) or three (green) bouts of control or drought conditions. This data includes both clitellate and aclitellate earthworms.

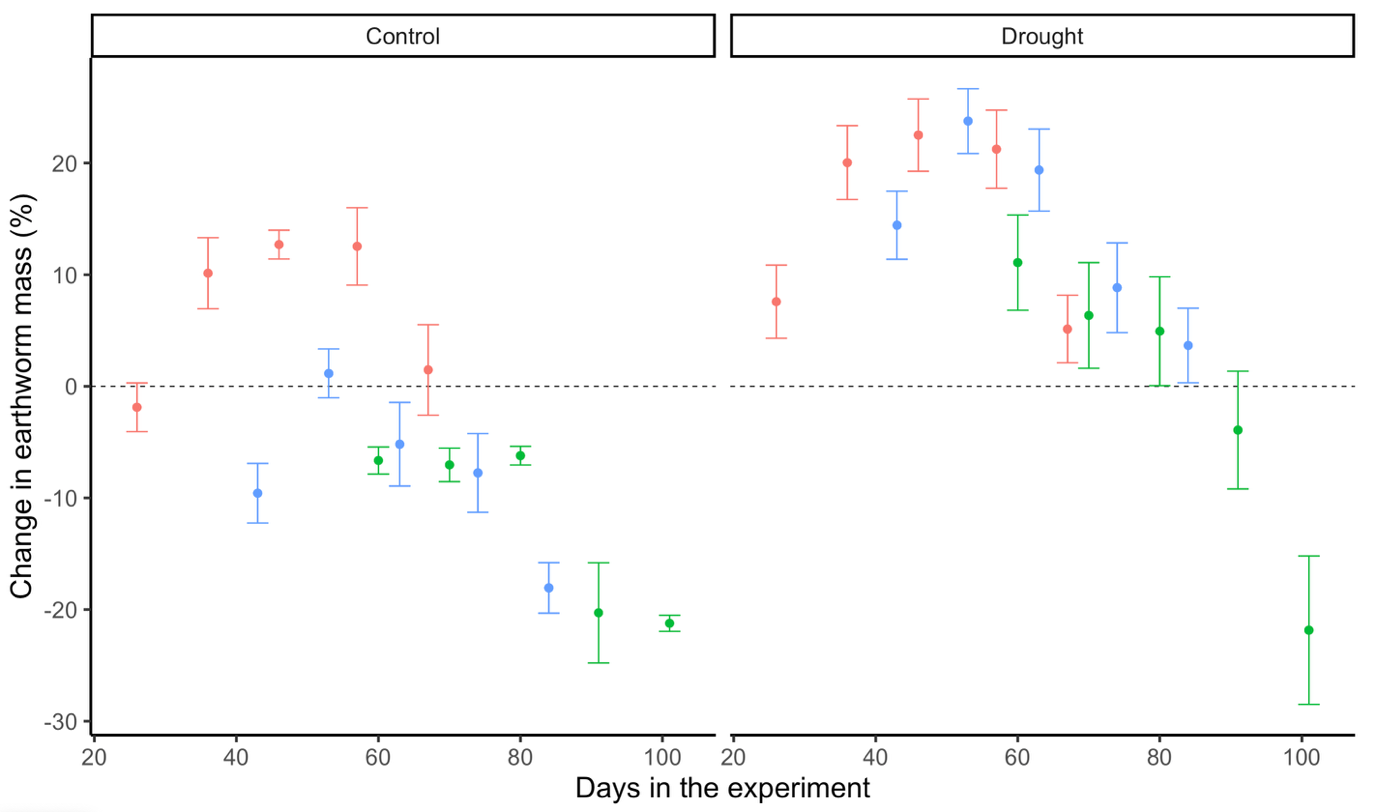
